# A new conceptual framework explaining spatial variation in soil nitrous oxide emissions

**DOI:** 10.1101/2023.11.27.568944

**Authors:** Ziliang Zhang, William C. Eddy, Emily R. Stuchiner, Evan H. DeLucia, Wendy H. Yang

## Abstract

Soil emissions of nitrous oxide (N_2_O), a potent greenhouse gas, contribute substantially to global warming from agriculture. Spatial variation in N_2_O emissions within agricultural fields leads to high uncertainty in the benefits of climate-smart agricultural practices. Here, we present a new conceptual framework explaining spatial variation in soil N_2_O emissions developed from high spatial resolution automated measurements of soil N_2_O emissions together with measurements of gross N_2_O fluxes and soil physicochemical properties in two separately managed maize fields in central Illinois, USA. We found that sub-field locations with consistently low N_2_O emissions had distinct biogeochemical properties compared to locations where high emissions occurred episodically, leading to spatial variation in which factors control N_2_O production rates. In the consistent N_2_O cold spots, soil nitrate (NO_3_^-^) and dissolved organic carbon (DOC) constrained N_2_O production irrespective of changes in soil moisture. In contrast, in the episodic N_2_O hot spots which had higher soil NO_3_^-^ and DOC availability, N_2_O production was stimulated by increases in soil moisture. These findings form the ‘cannon model’ which conceptualizes how sub-field scale variation in soil NO_3_^-^ and DOC determines where increases in soil moisture can trigger high soil N_2_O emissions within agricultural fields.

---

Nitrous oxide (N_2_O) currently accounts for nearly 6% of net radiative forcing in the Earth’s atmosphere^1^, with over half of rising atmospheric N_2_O concentrations attributed to agricultural activities^2, 3^. This potent greenhouse gas is produced in soil via microbially and chemically mediated processes which are highly sensitive to environmental conditions such as soil moisture, inorganic nitrogen availability, and temperature^4, 5^. The resulting spatial and temporal variability in soil N_2_O emissions causes large uncertainty in measurement and modeling of the global warming potential outcomes of agricultural management practices, including soil carbon sequestration practices that may inadvertently increase soil N_2_O emissions to offset climate change mitigation benefits^6, 7, 8, 9, 10^. Short-lived, exceptionally high soil N_2_O emissions that contribute disproportionately to annual N_2_O budgets can be triggered by events, such as rainfall, fertilization, and freeze-thaw^11, 12, 13, 14^. These N_2_O hot moments exhibit high spatial variation, even within agricultural fields with little topographic relief and in monoculture crop production^15, 16^. Due to methodological constraints in measuring soil N_2_O emissions, the lack of datasets that capture both high spatial and high temporal resolution has limited advances in understanding the drivers of spatial variation in temporal patterns of emissions.

## Insights from a high spatial and high temporal resolution N_2_O flux dataset

An unprecedented high spatial and high temporal resolution dataset revealed that a relatively flat agricultural field in commercial maize production harbored subfield locations that acted as consistent N_2_O cold spots versus episodic N_2_O hot spots (Figure 1ab). The dataset was generated from automated hourly net N_2_O flux measurements at 20 locations across a 4.6 ha area in the field (Figure S1). The cold spots had consistently below average net N_2_O fluxes compared to other locations in the field and did not experience N_2_O hot moments. This was indicated by both low mean relative difference (MRD) and low standard deviation relative difference (SDRD) of mean daily N_2_O fluxes over the 2021 growing season (Figure 1ab). In contrast, the hot spots had both high MRD and high SDRD (Figure 1ab), which reflects the contribution of infrequent N_2_O hot moments to both high mean fluxes and high variation in fluxes over the growing season. The absence of consistent N_2_O hot spots exhibiting high MRD and low SDRD suggests that high net N_2_O fluxes occurred only when episodically triggered by the occurrence of favorable environmental conditions.

**Figure 1.**
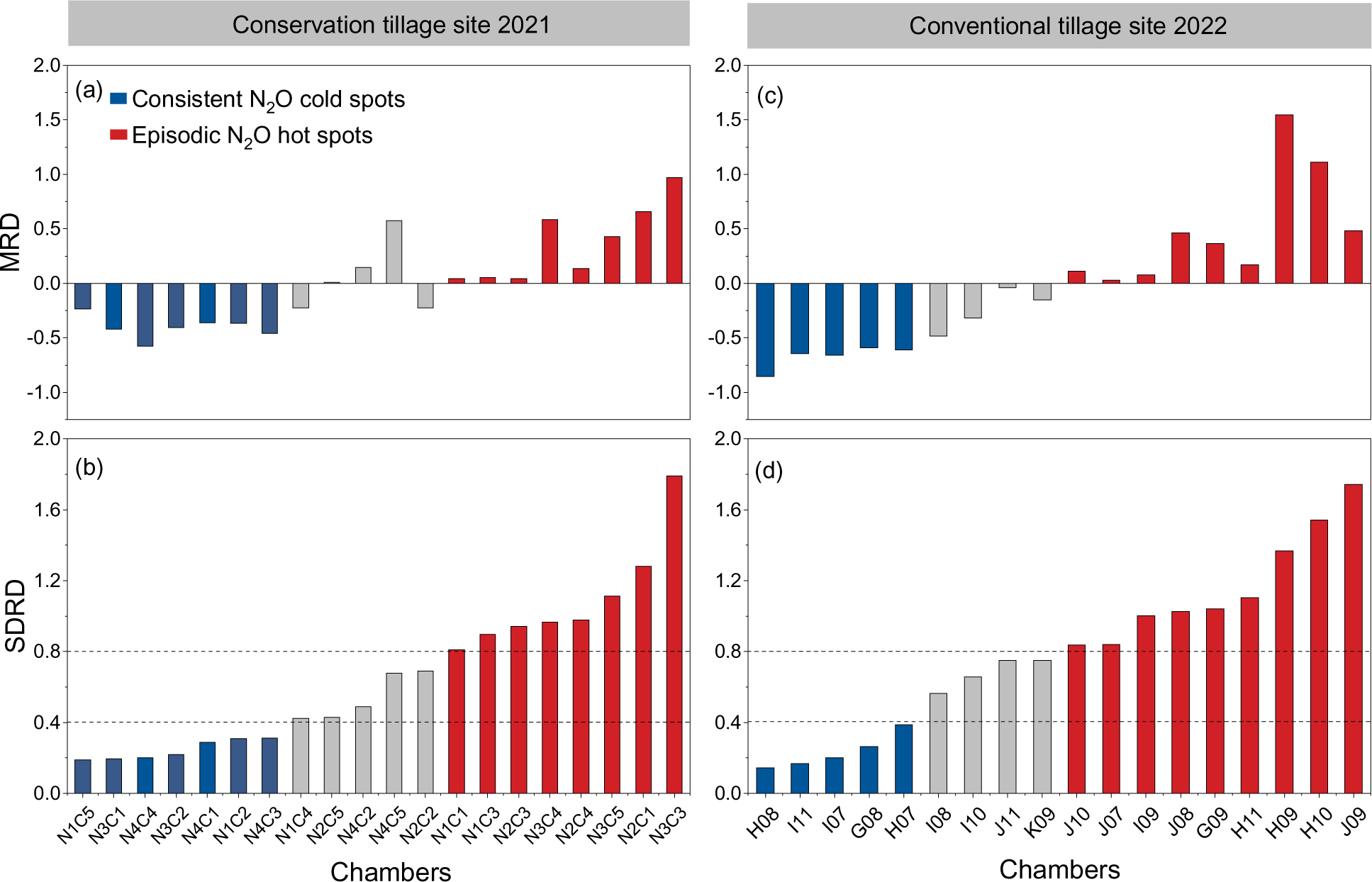
(a) Mean relative difference (MRD) and (b) standard deviation relative difference (SDRD) of daily net nitrous oxide (N_2_O) fluxes averaged from hourly flux measurements from June 2021 to October 2021 at 20 autochamber locations in a maize field under conservation tillage. (c) MRD and (d) SDRD of net N_2_O fluxes measured using manual static flux chambers on 12 sampling dates from May 2022 to October 2022 at 18 locations in a 50 m x 50 m grid in a maize field under conventional tillage. For 2021, sampling locations are identified by sampling node (N1-N4) and autochamber within the node (C1-C5). For 2022, sampling locations are identified by grid position. Blue bars indicate sampling locations identified as consistent N_2_O cold spots based on MRD below zero and SDRD below 0.4; red bars indicate sampling locations identified as episodic N_2_O hot spots based on MRD above zero and SDRD above 0.8; gray bars indicate sampling locations considered intermediate in MRD and SDRD. The dotted lines mark the SDRD thresholds of 0.4 and 0.8 for identifying the cold and hot spots.

High net N_2_O fluxes were caused by stimulation of N_2_O production, largely from denitrification. Spatial and temporal variation in gross N_2_O production rates spanned an order of magnitude greater range than gross N_2_O consumption rates (Figure 2ab, de), based on monthly *in situ* ^15^N_2_O pool dilution measurements at all autochamber locations over the growing season. As such, patterns in net N_2_O fluxes mirrored gross N_2_O production (Figure 2c, f), with a strong positive correlation between net N_2_O fluxes and gross N_2_O production rates (R^2^ = 0.90, N = 100, *P* < 0.001; Figure S2a). The importance of denitrification as an N_2_O source process has been documented in other agricultural systems^17, 18^. This was also demonstrated in our field site using ^15^NH_4_^+^ versus ^15^NO_3_^-^ tracers to partition N_2_O production from nitrification versus denitrification in 4-hour laboratory incubations of soil samples collected near each autochamber (Figure S3a). Furthermore, we found that denitrification was a more important N_2_O source in the episodic N_2_O hot spots where high N_2_O production occurred (Figure S3a). These findings together suggest that understanding controls on N_2_O production via denitrification in the environment will improve predictions of spatiotemporal variation in net N_2_O fluxes.

**Figure 2.**
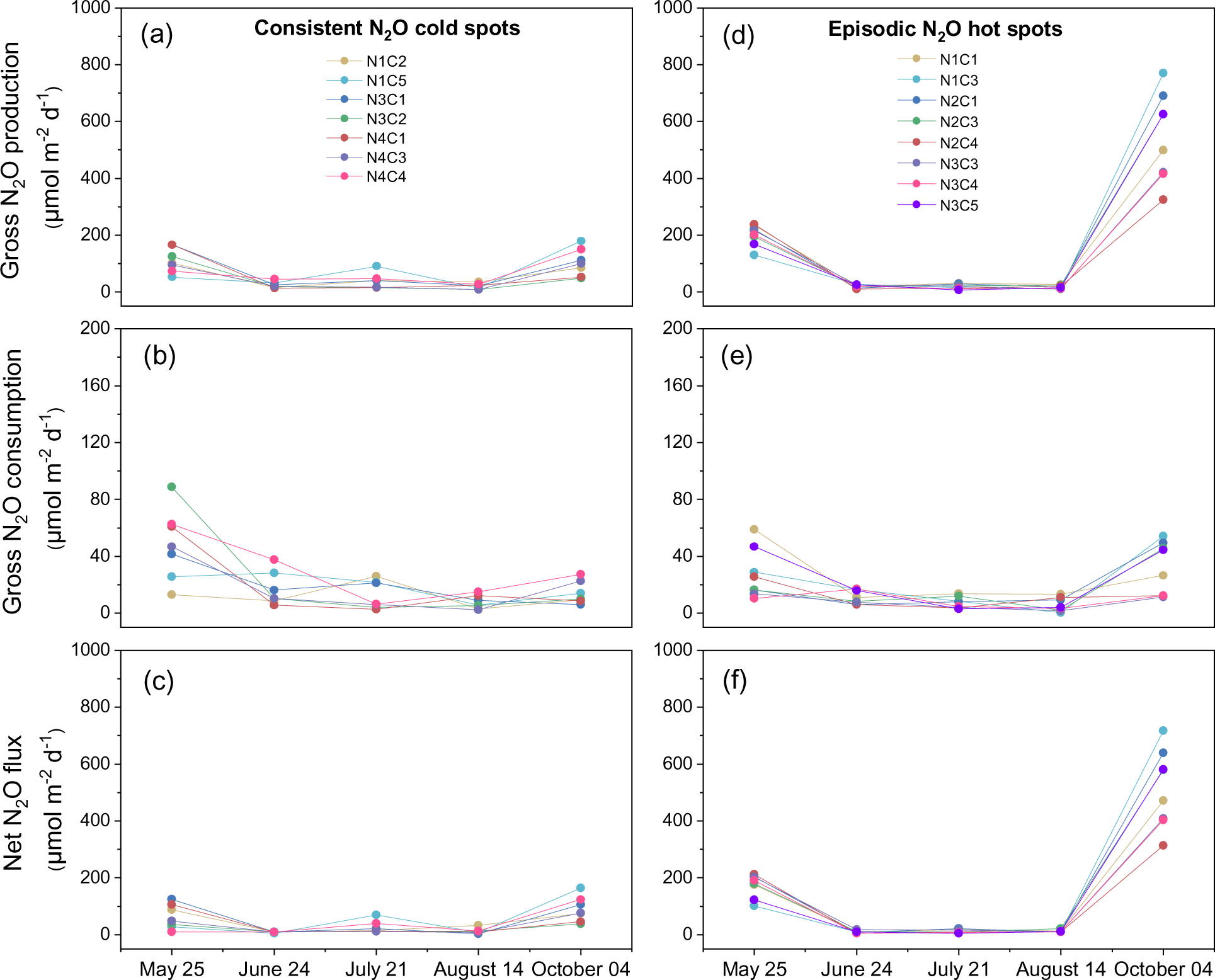
Gross nitrous oxide (N_2_O) production rates, gross N_2_O consumption rates, and net N_2_O fluxes measured monthly in consistent N_2_O cold spots (a, b, c, respectively) and episodic N_2_O hot spots (d, e, f, respectively) in the conservation tillage site in maize production in 2021. Colors represent different sampling locations, which are identified by sampling node (N1-N4) and autochamber within the node (C1-C5).

## Dominant controls on N_2_O production vary spatially within fields

The dominant drivers of gross N_2_O production rates differed between consistent N_2_O cold spots versus episodic N_2_O hot spots (Figure 3ab, S4ab). Structural equation modeling revealed that, in cold spots, nitrate (NO_3_^-^) and dissolved organic carbon (DOC) concentrations had major positive, direct effects on gross N_2_O production whereas soil water-filled pore space (WFPS) had only a minor indirect effect (Figure 3a). By comparison, in hot spots, WFPS and iron (Fe) redox status—indices of anoxia in bulk soil and soil microsites, respectively—had major positive direct effects on gross N_2_O production whereas NO_3_^-^ had a minor direct effect and DOC had no effect (Figure 3b). Soil moisture, NO_3_^-^, and DOC are well-known controls on denitrification, an anaerobic microbial process by which NO_3_^-^ is reduced by organic C^4, 19^. Yet, the differing hierarchal importance of these predictor variables in consistent N_2_O cold spots versus episodic N_2_O hot spots has not previously been recognized.

**Figure 3.**
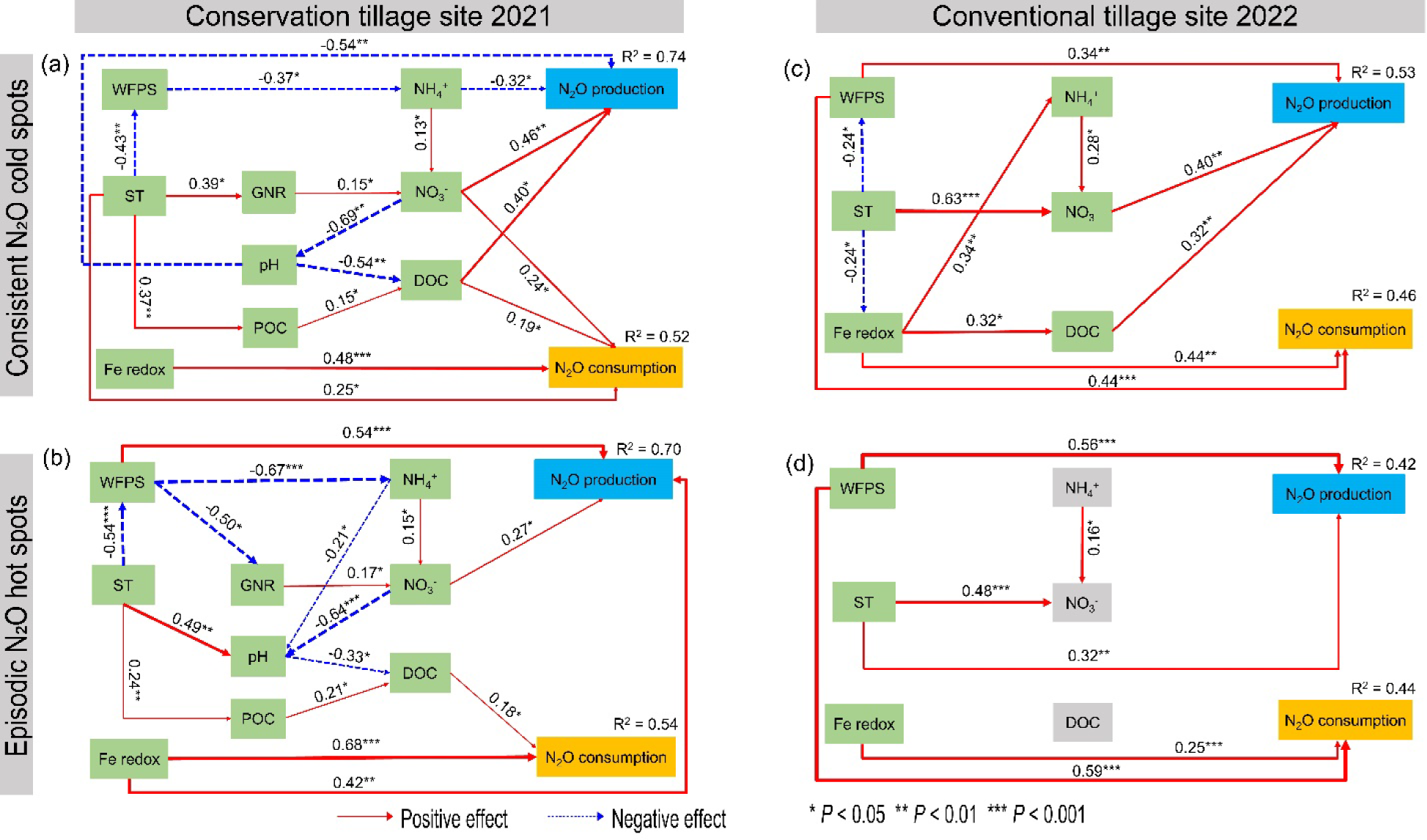
Partial least squares structural equation model showing effects of soil variables on gross nitrous oxide (N_2_O) production and consumption (a, b) measured monthly over the 2021 growing season in consistent N_2_O cold spots and episodic N_2_O hot spots, respectively, at the conservation tillage site and (c, d) measured at 12 time points over the 2022 growing season in cold versus hot spots, respectively, at the conventional tillage site. Arrow heads indicate the hypothesized direction of causation, and arrow width is proportional to the strength of the relationship. Solid red arrows indicate positive effects; the dashed blue arrows indicate negative effects. Numbers by the arrows are the standardized path coefficients with * *p* <0.05, ** *p* <0.01, *** *p* <0.001. Soil variables were measured at both sites include: dissolved organic carbon (DOC), iron redox status (Fe redox), microbial biomass carbon, soil ammonium (NH_4_^+^), soil nitrate (NO_3_^-^), soil pH, soil temperature (ST), and water-filled pore space (WFPS). Gross mineralization rate, gross nitrification rate (GNR), and particulate organic carbon (POC) were measured only at the conservation tillage site. Insignificant effects are not shown.

**Figure 4.**
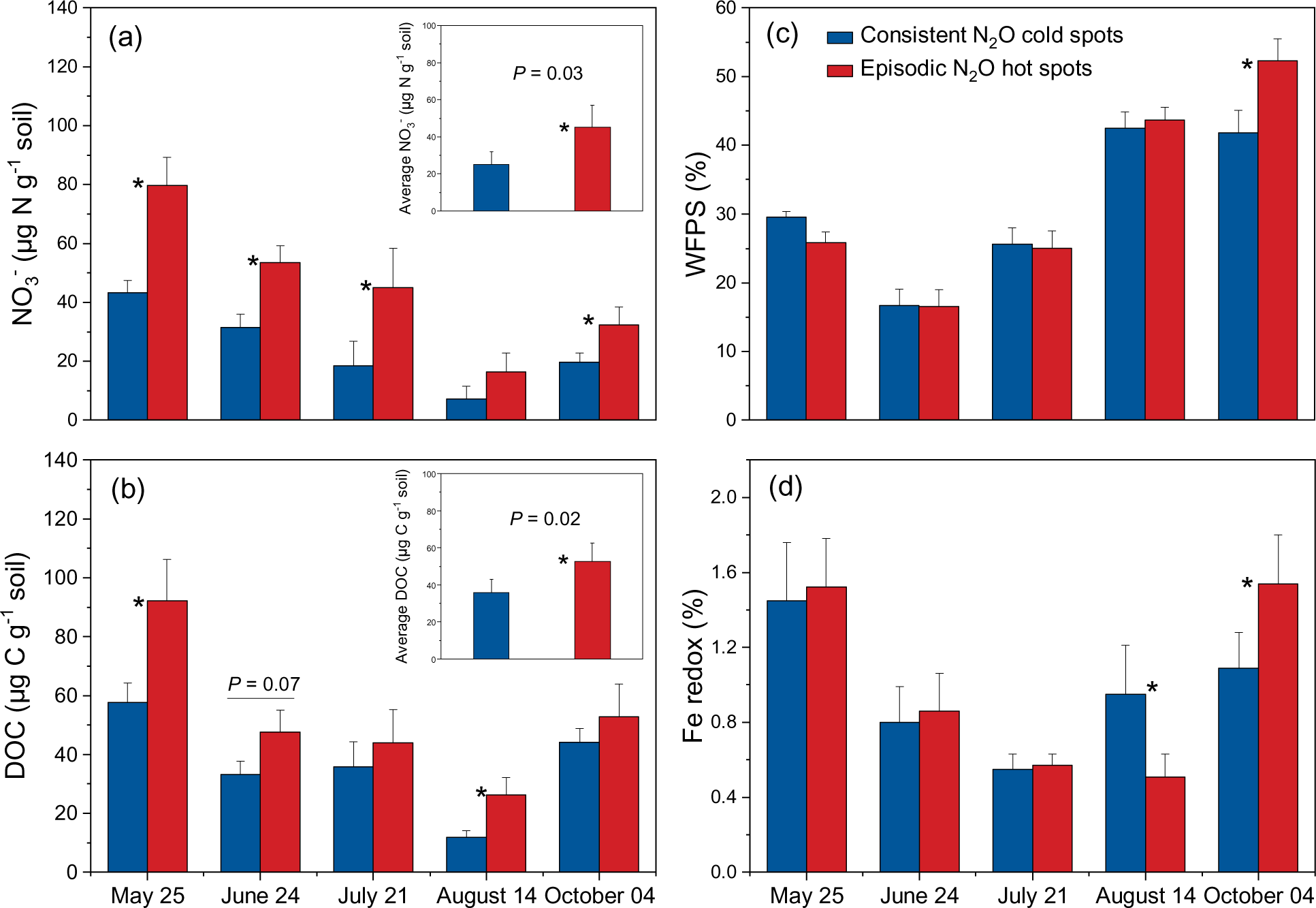
(a) Soil nitrate (NO_3_^-^) concentrations, (b) dissolved organic carbon (DOC) concentrations, (c) water-filled pore space (WFPS), and (d) iron (Fe) redox status (Fe(II) percentage of total 0.5 N HCl-extractable Fe pool, an index of anaerobic soil microsites) in consistent nitrous oxide (N_2_O) cold spots (blue bars) versus episodic N_2_O hot spots (red bars) measured monthly over the 2021 growing season at the conservation tillage site. Planting occurred on May 1, 2021, and post-plant fertilization with UAN 32% at a rate of 202 kg N ha^-1^ occurred on May 7, 2021. Asterisks denote significant differences between cold versus hot spots at *P* < 0.05 level. Error bars represent standard errors (n=7 and 8 for cold spots and hot spots, respectively).

The spatial variation in dominant controls on soil N_2_O production was corroborated using a separately managed commercial maize field under conventional tillage in central Illinois, USA in the 2022 growing season. To capture a greater range in soil conditions, 18 sampling locations were distributed in a 50 m × 50 m grid within a 5 ha area and 12 soil sampling dates were timed to represent conditions just after rain events or during periods with little rainfall (Figure S5b). Similar to the conservation tillage site, the sampling locations at this site could be categorized as consistent N_2_O cold spots or episodic N_2_O hot spots (Figure 1cd). Net N_2_O flux patterns again mirrored patterns in gross N_2_O production (Figure S2c, S6). At this site, gross N_2_O production rates were also dominantly controlled either by NO_3_^-^ and DOC or by WFPS in the cold spots versus hot spots, respectively (Figure 3cd, S4cd). The consistency in results between two differently managed maize fields in different growing seasons supports the generalizability of these results.

Lower soil concentrations of NO_3_^-^ and DOC in consistent N_2_O cold spots compared to episodic N_2_O hot spots throughout the growing season at both sites suggests that substrate availability constrained denitrification rates in the cold spots (Figure 4ab, S7ab, Tables S1-S4). Soil WFPS and Fe redox status generally did not differ between cold and hot spots at either site (Figure 4cd, S7cd, Tables S1-S4). This discounts the possibility that higher soil O_2_ suppressed denitrification in the cold spots. Instead, soil N_2_O production in the episodic N_2_O hot spots could be stimulated by increases in soil moisture due to sufficient NO_3_^-^ and DOC availability to support high denitrification rates. The same increases in soil moisture in the cold spots could not stimulate N_2_O production due to more limited NO_3_^-^ and DOC availability. The difference in dominant controls on N_2_O production between cold and hot spots is therefore ultimately determined by soil NO_3_^-^ and DOC availability, as conceptualized in the ‘cannon model’ (Figure 5).

**Figure 5.**
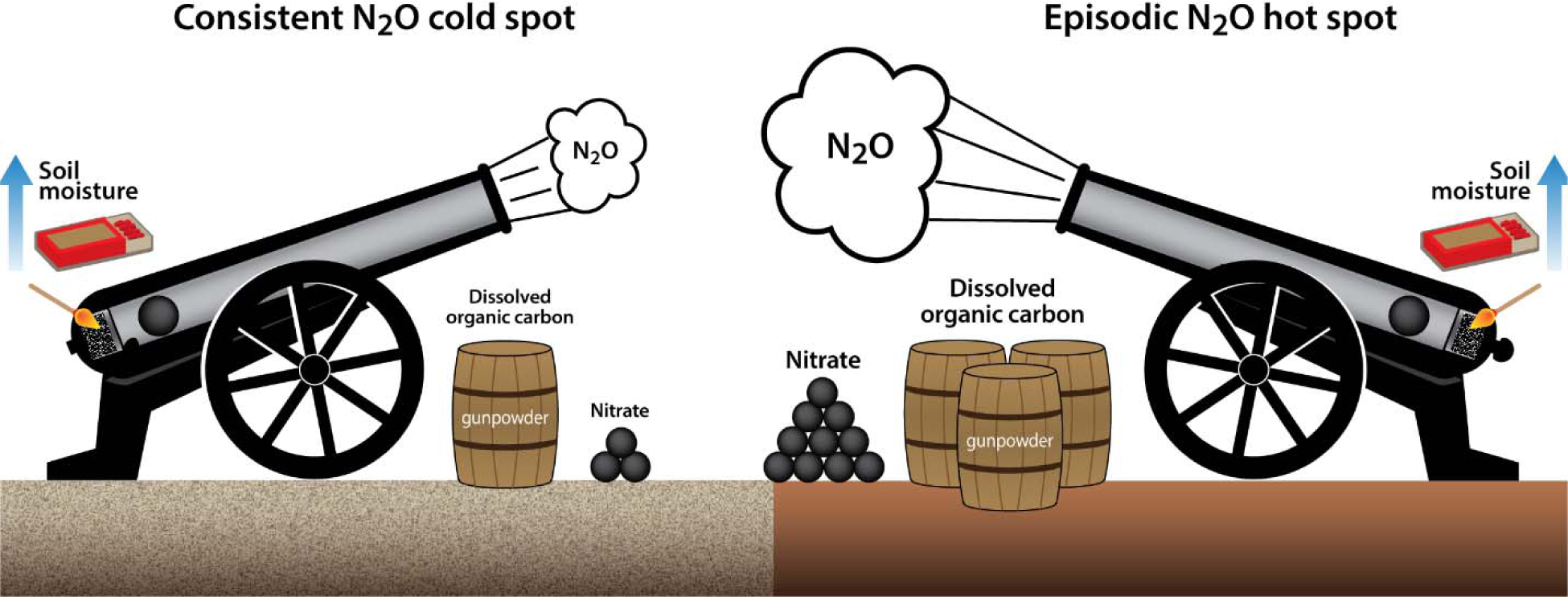
The ‘cannon model’ conceptualizes the different hierarchical controls on soil nitrous oxide (N_2_O) production in consistent N_2_O cold spots versus episodic N_2_O hot spots during the growing season. Nitrate (NO_3_^-^) is the substrate that is reduced to N_2_O by denitrifying microbes using electrons donated from organic carbon when high soil moisture creates anoxic soil conditions conducive for the anaerobic process of denitrification. In locations with greater soil NO_3_^-^ and dissolved organic carbon (DOC) availability, soil moisture is the dominant variable controlling variation in N_2_O production rates, with increases in soil moisture causing these locations to experience an N_2_O hot moment. In contrast, in locations with more limited availability of soil NO_3_^-^ and DOC, soil N_2_O emissions are constrained by low N_2_O production rates that vary primarily based on soil NO_3_^-^ and DOC concentrations. This is akin to how lighting the ignition chamber of a cannon (increasing soil moisture) can lead to repeated firing of the cannon (an N_2_O hot moment) only when there is sufficient cannonballs (NO_3_^-^) and gunpowder (DOC).

## Potential drivers of spatial variation in soil NO_3_^-^ and DOC

The seasonal pattern in soil NO_3_^-^ concentrations in consistent N_2_O cold spots versus episodic N_2_O hot spots suggests that greater rates of NO_3_^-^ consumption, by soil microbes and plants, contributed to the lower soil NO_3_^-^ in the cold spots (Figure 4a, S7a). At the conventionally tilled site in 2022 when soil sampling was more frequent, soil NO_3_^-^ was comparably low across the field until six days after spring fertilization when soil NO_3_^-^ was comparably elevated across the field. Lower soil NO_3_^-^ in the cold spots was first detected at 11 days post-planting (16 days post-fertilization) and persisted through the remainder of the growing season, although the difference became statistically not significant in the late growing season as soil NO_3_^-^ decreased overall. Given that little rainfall preceded the spatial variation in soil NO_3_^-^ developing (Figure S1, S5b), greater microbial consumption of NO_3_^-^ in the cold spots presumably led to the lower soil NO_3_^-^ at least initially. The greatest differences in soil NO_3_^-^ between cold and hot spots occurred in the mid-growing season, following side-dress fertilization application synchronized with high plant N demand (Figure S7a). This suggests that greater plant N uptake of fertilizer N in the cold spots may have also played a role in creating the spatial patterns in soil NO_3_^-^. Determining the mechanisms leading to greater plant and microbial N consumption in the cold spots could not only improve predictions of spatial variation in soil N_2_O emissions but also guide the development of novel strategies for mitigating emissions.

Lower DOC concentrations in consistent N_2_O cold spots may be caused by more limited supply of DOC from the soil organic carbon (SOC) pool. At both sites, bulk SOC concentrations were lower in the cold spots compared to the episodic N_2_O hot spots, with both lower particulate organic carbon (POC) and mineral-associated organic carbon (MAOC) concentrations contributing to this pattern (Table 1). Sub-field scale spatial variation in denitrification potential has been linked to variation in POC^20^. Although DOC can be derived from both the POC and MAOC fractions of SOC, C turnover rates in MAOC are generally slower due to chemical protection of OC via adsorption to soil minerals^21, 22^. In addition to POC serving as an important source of DOC (Figure S8), POC-derived DOC stimulates denitrification more than DOC derived from MAOC^23^. This suggests that both higher quantity and quality of POC-derived DOC may lead to N_2_O production in the hot spots not being limited by OC. Given that POC consists of partially decomposed plant material^22, 24^, understanding controls on spatial variation in aboveground plant residues and belowground plant productivity can potentially inform predictions of POC that may underlie the spatial patterns in DOC.

**Table 1.**
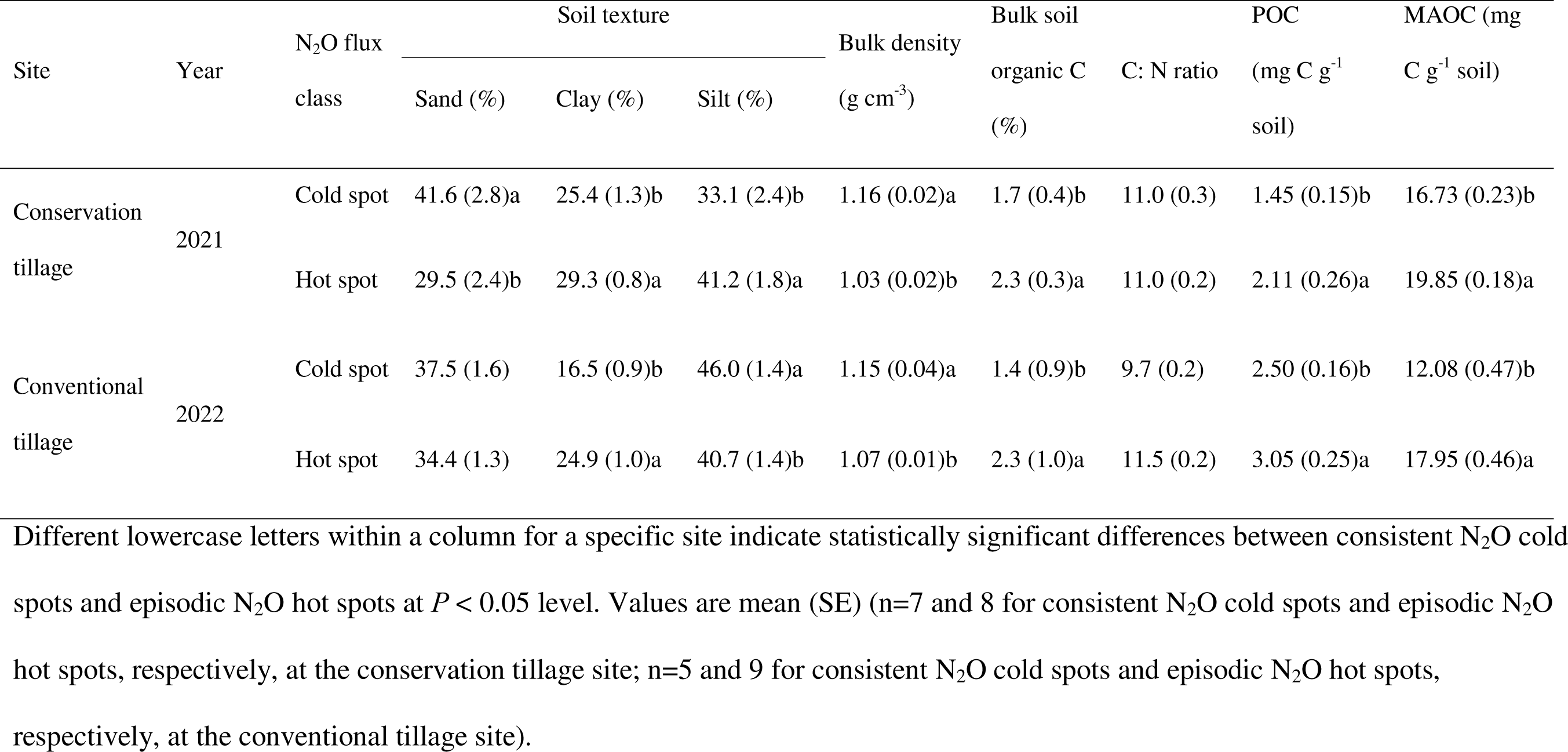
Soil physical and chemical properties by N_2_O flux class at the two study sites in maize-soybean rotations, with maize grown in the study year.

Soil texture differences may also contribute to the differences in soil NO_3_^-^ and DOC availability between consistent N_2_O cold spots and episodic N_2_O hot spots. At both field sites, the cold spots had silt loam soils with higher sand content and lower clay content than silty clay loam soils in the hot spots (Table 1). Positive relationships between SDRD of net N_2_O fluxes (a measure of temporal variability) and the ratio of clay content to sand content at both sites suggests that the potential for high soil N_2_O emissions to occur increases as clay content increases and sand content decreases (R^2^ = 0.45-0.50, *P* < 0.001, Figure S9).

Drainage is greater in sandier soils such that the greater leaching of dissolved soil constituents can lead lower soil NO_3_^-^ concentrations^25, 26^. At the same time, DOC can adsorb to clay minerals, resulting in greater retention of DOC with greater clay content^27, 28, 29^. Soil texture can also influence plant and microbial access to nutrients^30, 31^ and SOC dynamics^32, 33^ to contribute to spatial patterns in soil NO_3_^-^ and DOC. Sub-field variation in soil texture may result from the accumulated impact of subtle, long-term patterns in surface hydrology that transported clay into localized micro-depressional areas now characterized by higher clay content^34^. Soil texture therefore represents a relatively static soil property that could be used to predict the locations of consistent N_2_O cold spots and episodic N_2_O hot spots within agricultural fields.

## A new conceptual framework for predicting soil N_2_O emissions

Here, we present a new conceptual framework that advances the prediction of spatial variation in soil N_2_O emissions within agricultural fields (Figure 5). Prior conceptual frameworks have predicted N_2_O hot spots based on topographic relief at landscape to watershed scales^11, 35^, which could not explain high spatial variation in N_2_O emissions observed within fields with little topography^15, 16^. In those frameworks, DOC and NO_3_^-^ moving downslope with water leads to convergence of all three denitrification controlling factors in foot slopes or riparian areas where N_2_O hot spots are often observed ^36, 37, 38, 39^. The ‘cannon model’ can be applied both within and beyond the context of topographic relief by generally conceptualizing how high soil N_2_O emissions can be triggered by increases in soil moisture only in locations with sufficiently high availability of soil NO_3_^-^ and DOC (Figure 5).

The ‘cannon model’ provides a new framework to guide measurement, modeling, and mitigation of agricultural soil N_2_O emissions. First, the model presents that sub-field scale variation in soil NO_3_^-^ and DOC determines spatial patterns in consistent N_2_O cold spots versus episodic N_2_O hot spots. This can inform efforts to measure soil N_2_O emissions and scale up the measurements to accurately estimate ecosystem-scale N_2_O budgets. Second, the model illustrates the different hierarchical importance of soil moisture, NO_3_^-^ and DOC in controlling N_2_O production rates in cold versus hot spots. This suggests that spatially explicit ecosystem models can more accurately predict soil N_2_O emissions by representing spatially varying dominant controls on N_2_O production. Third, the model highlights the role of higher soil NO_3_^-^ and DOC availability in creating potential N_2_O hot spots. Understanding the drivers of spatial variation in soil NO_3_^-^ and DOC, which may be related to soil texture, is therefore the key to developing precision agricultural practices that target reductions in N_2_O emissions from hot spots that disproportionately contribute to field-scale N_2_O budgets^12, 40^. This also suggests another way in which climate-smart agricultural practices aimed at increasing SOC may inadvertently increase soil N_2_O emissions^6, 7, 8^, by increasing DOC and soil NO_3_^-^ derived from soil organic matter to turn cold spots into hot spots. Overall, this conceptual breakthrough in understanding controls on spatial variation in soil N_2_O emissions holds promise for guiding future efforts to reduce uncertainty in and effectively mitigate agricultural soil N_2_O emissions.

## Methods

This study was conducted in two separately managed commercial agricultural fields in maize production located in Champaign County, Illinois, USA. One field managed with conservation tillage was sampled in the 2021 growing season (hereafter referred to as the “conservation tillage site”), and the other field managed with conventional tillage was sampled in the 2022 growing seasons (hereafter referred to as the “conventional tillage site”). Detailed site descriptions and management activities are provided in the Supplementary methods.

To capture spatial and temporal variability in soil N_2_O emissions at the field scale, net soil-atmosphere fluxes of N_2_O were measured hourly using autochambers at 20 locations within the conservation tillage site. The autochambers were distributed in four sampling nodes within a 5 ha area of the field with high spatial variation in net N_2_O flux patterns observed in the field the prior year using 50 m x 50 m grid sampling with weekly to monthly manual flux measurements (Nakian Kim, unpublished data). At each node, five LI-COR autochambers were radially installed at 12 m distance from a N_2_O gas analyzer (LI-7820, LI-COR Biosciences, Lincoln, NE, USA) that sequentially measured hourly net soil-atmosphere N_2_O fluxes from each autochamber continuously starting in June 2021. Time stability (TS) analysis was employed according to Ashiq et al.^41^ to identify consistent N_2_O cold spots and episodic N_2_O hot spots. Chamber locations with low mean relative difference (MRD, negative) and low standard deviation of relative difference (SDRD, < 0.4) were classified as consistent N_2_O cold spots, and chamber locations with high SDRD (≥ 0.8) were classified as episodic N_2_O hot spots. To validate the SDRD thresholds used for this classification, a grouping analysis was performed using the spatial statistics tools in ArcMap 10.8.1 (ESRI, CA, USA). More details about the time stability analyses are provided in the Supplementary methods.

We used the ^15^N_2_O pool dilution technique to measure gross N_2_O fluxes (i.e., gross N_2_O production and consumption) in the field over the growing season (May to October). At the conservation tillage site in 2021, these measurements were conducted monthly adjacent to (0.5 m away from) the 20 autochamber locations. At the conventional tillage site in 2022, these measurements were conducted on 12 sampling dates at 18 locations in a 50 m x 50 m grid. We performed the measurements over 45 minutes using static flux chambers as described by Krichels et al.^42^. Details are provided in the Supplementary materials. After the last gas sample was collected from a chamber, we measured soil temperature and soil volumetric water content at 0-10 cm depth in the chamber footprint using an Acorn Temp 5 meter (Oakton Instruments, Vernon Hills, IL, USA) and a hand-held moisture meter (HH2 moisture meter; Delta-T Devices Ltd, Cambridge, UK), respectively. Gas samples were analyzed for CO_2_, N_2_O, and SF_6_ concentrations on a gas chromatograph (GC, Shimadzu Scientific Instruments, Columbia, MD, USA) equipped with a thermal conductivity detector (TCD) and an electron capture detector (ECD). The gas samples were also analyzed for ^15^N isotopic composition of N_2_O on a IsoPrime 100 isotope ratio mass spectrometer (IRMS) interfaced to a trace gas preconcentration unit (Isoprime Ltd., Cheadle Hulme, UK) and a GX-271 autosampler (Gilson, Inc., Middleton, WI, USA). Gross N_2_O production and consumption rates were calculated from the change in ^14^N_2_O, ^15^N_2_O, and SF_6_ concentrations over time using the pool dilution model as described by Yang et al.^43^. Net N_2_O fluxes from the manual chamber measurements were calculated from the exponential change in N_2_O concentration over time^44^. Net N_2_O flux was considered to be zero when the relationship between N_2_O concentration and time was not significant (*P* > 0.05).

Immediately after each ^15^N_2_O pool dilution measurement was completed in the field, a soil sample from the chamber footprint was collected to partition N_2_O source processes and measure soil properties potentially controlling soil N_2_O dynamics. Two soil samples (0-20 cm depth) were collected from the chamber footprint using a soil auger (5 cm diameter), composited, and then split for the various assays. On the same day as soil collection, we performed 2 M KCl and 0.5 N HCl extractions on subsamples of the composited soil samples to characterize soil inorganic N availability and iron (Fe) redox status, respectively, near the chamber locations. Iron redox status, as a proxy for the abundance of anaerobic soil microsites, was calculated as the percentage of the total acid-extractable Fe pool accounted for by Fe(II). Within 24 h of soil collection from the field, we performed 4 h ^15^N pool dilution measurements in the laboratory to quantify gross rates of mineralization (GMR) and nitrification (GNR) using ^15^NH_4_Cl and K^15^NO_3_, respectively^45, 46^. The added ^15^N also served as tracers to estimate N_2_O production from nitrification and denitrification based on ^15^N_2_O produced in soils receiving ^15^NH_4_^+^ versus ^15^NO_3_^-^, respectively^42^. Within 48 h after soil collection, fresh soil subsamples were extracted in a 3:1 ratio of deionized water to dry soil equivalent mass for determination of dissolved organic carbon (DOC) concentrations and in 0.5 M K_2_SO_4_ for determination of soil microbial biomass C (MBC) by direct chloroform extraction as described by Setia et al. (2012)^47^. Soil gravimetric water content (GWC) was measured by oven-drying 10-g subsample at 105 °C for 24 h. Air-dried soil subsamples were used for measurements soil pH and concentrations of soil organic C (SOC) and total (TN) as well as for soil physical fractionation. Concentrations of particulate organic carbon (POC) and mineral-associated organic carbon (MAOC) were determined after size fractionation using the method modified from Cotrufo et al.^21^ and Zhang et al.^48^. Given that non-significant temporal variation of SOC concentration was detected over the growing season of 2021 at the conservation tillage site, concentrations of SOC, POC, and MAOC were only quantified on two sampling dates (one date representing early growing season and another one representing late growing season) over the growing season of 2022 at the conventional tillage site. Details about sample analyses are reported in the Supplementary methods.

Soil texture and bulk density at both sites were measured from soil samples collected at each chamber location for the ^15^N_2_O pool dilution measurement at the end of the growing season. Two intact soil cores (0-20 cm depth) were taken from each chamber location using a 5 cm diameter stainless-steel quantitative soil corer. One soil core was used to measure bulk density that was calculated as the dry soil weight by dividing the volume of the core after removing visible rocks and plant materials^15^. The other soil core was used to measure soil texture that was determined using a hydrometer after dispersion with 5% sodium hexametaphosphate solution as described by Gavlak et al.^49^. Soil water-filled pore space (WFPS) was calculated using GWC and bulk density, assuming a soil particle density (PD) of 2.65 g cm^-3^ ^50^.

All statistical analyses were performed using R 4.0.4 (R Core Team, 2021). All data and residuals were tested for normality before the data analysis. Differences were considered significant at the *P* < 0.05 level. The data for each field site were analyzed separately. We used repeated measures ANOVA to compare soil properties and N_2_O fluxes between consistent N_2_O cold spots and episodic N_2_O hot spots (between-subjects factor) with sampling date as a repeated factor (within-subjects factor). For a specific sampling date, independent t-tests were used to assess the difference in all variables between N_2_O flux classes. For soil properties measured only once (i.e., soil texture and bulk density), differences between N_2_O flux classes were also analyzed using independent t-tests. A Pearson correlation analysis was conducted to identify statistically significant correlations between soil physicochemical properties and gross N_2_O production and consumption rates using the “corrplot” R package^51^. Partial least squares structural equation models (PLS-SEM) were used to determine the direct and indirect effects of soil variables on gross N_2_O production and consumption using the “plspm” R package^52^. The results from the Pearson correlation analysis served as the hypothetical base for the initial PLS-SEM model. The PLS-SEM analyses were conducted separately for consistent N_2_O cold spots and episodic N_2_O hot spots.

## Acknowledgements

We appreciate assistance from DoKyoung Lee, Nakian Kim, and Adam von Haden in selecting the sampling locations, Adam von Haden in setting up the autochamber equipment, Ayesha Ahmed in maintaining the autochamber measurements, and Allison Cook, Ingrid Holstrom, Jessica Mulcrone, Haley Ware, and Chloe Yates in autochamber installation and removal at the conservation tillage site. We appreciate Chloe Yates, Samantha Davis, Ava Bernacchi, and Neiman Shivers assisting in the field and laboratory with the isotope-based measurements and soil analyses. This study was supported by the U.S. Department of Energy ARPA-E SMARTFARM program under Award Number DE-AR0001382.

## Author contributions

Z.Z, W.C.E., E.H.D., and W.H.Y. designed the study; Z.Z. and W.C.E. led data collection, processing, and analysis; Z.Z., W.C.E., W.H.Y., E.S., and E.H.D. interpreted the data; W.C.E. developed the cannon model; Z.Z. and W.H.Y. wrote the manuscript with contributions from W.C.E., E.H.D., and E.S.

## Notes

### Competing Interest Statement

The authors have declared no competing interest.

