## Supplementary material for "A new conceptual framework explaining spatial variation in soil nitrous oxide emissions"

**Supplementary Information**

**1. Supplementary Methods**

**2. Supplementary Tables**

**3. Supplementary Figures**

**4. Supplementary References**

**1. Supplementary Methods**

**Site description**

This study was conducted in two separately managed commercial agricultural fields located in Champaign County, Illinois, USA. One field located near Bondville, IL is cultivated in maize-soy rotations with conservation tillage and was planted with maize (*Zea mays*) during the 2021 growing season (hereafter referred to as the “conservation tillage site”). The soil in this field is roughly evenly split among Drummer silty clay loam, Flanagan silt loam, and Catlin silt loam (USDA-NRCS, 2022). The other field located near Villa Grove, IL is cultivated in maize-maize-soy rotations with conventional tillage and was planted with maize during the 2021 and 2022 growing seasons (hereafter referred to as the “conventional tillage site”). The soil in the field is roughly 70% Drummer silty clay loam and 30% Millbrook silt loam (USDA-NRCS, 2022).

At the conservation tillage site, *Z. mays* was planted in early May of 2021 and followed with fertilizer application (202 kg N ha^−1^ as 32% urea ammonium nitrate (UAN)) and was harvested at the end of September. Following harvesting in late September, fall fertilization with monoammonium phosphate (MAP, 11-52-0) at 21 kg N ha^−1^ and 100 kg P ha^−1^ and potash at 62 kg K ha^−1^ occurred in early October. At the conventional tillage site, *Z. mays* was planted on May 12, 2022 after fertilizer application (19.7 kg N ha^-1^ and 91.3 kg P ha^-1^ as MAP, 134.5 kg N ha^−1^ as anhydrous ammonia, and 58.9 kg K ha^-1^ as potash) on May 7, 2022. Fertilizer side dressing with 90.2 kg N ha^−1^ as 32% UAN and 13.2 kg N ha^-1^ as ammonium thiosulfate occurred on June 11, 2022. *Z. mays* was harvested on October 28, 2022.

During the period of 1989-2012, the mean annual air temperature in Champaign County was 10 °C, with a maximum monthly mean temperature of 24.4 °C in July and a minimum of -5.5 °C in January. The mean annual precipitation during that period was 1008 mm (Illinois-Climate-Network, 2017). Daily air temperature and precipitation for the study periods (May to October of 2021 and 2022) were obtained from a climate station near the field sites which reports data to the Midwestern Regional Climate Center (Figure S5).

**Identification of spatiotemporal patterns in soil N_2_O emissions**

Time stability (TS) analysis was employed according to Ashiq et al.^1^ to identify consistent N_2_O cold spots and episodic N_2_O hot spots. A positive mean relative difference (MRD) value represents higher N_2_O emissions than the spatial average of the N_2_O emissions across all chamber locations over the growing season, while a negative MRD value represents less emissions than the spatial average emissions. A small value of standard deviation of relative difference (SDRD) indicates low temporal variation in emission, while a large SDRD value indicates high temporal variation in emission. Chamber locations with low MRD (negative) and low SDRD (< 0.4) were classified as consistent N_2_O cold spots, and chamber locations with high SDRD (≥ 0.8) were classified as episodic N_2_O hot spots. For the conservation tillage site, these analyses were performed on mean daily net N_2_O fluxes calculated from hourly measurements at 20 autochamber locations over the 2021 growing season. For the conventional tillage site, these analyses were performed on manual chamber-based measurements of net N_2_O fluxes at 18 locations on 12 sampling dates over the 2022 growing season. Chamber locations that did not fall into either of those spot classes were excluded from further analyses; this represented 25% of chamber locations at both sites. To validate the SDRD thresholds used for this classification, a grouping analysis was performed using the spatial statistics tools in ArcMap 10.8.1 (ESRI, CA, USA). High R^2^ values computed for SDRD indicates confidence in effectively separating different groups of temporal stability in net N_2_O fluxes^1^.

**Gross N_2_O flux measurements**

We used the ^15^N_2_O pool dilution technique to measure gross N_2_O fluxes (i.e., gross N_2_O production and consumption) in the field over the growing season (May to October). Briefly, we injected 10 ml of spiking gas consisting of 9.3 ppm sulfurhexafluoride (SF_6_) and 18.5 ppm N_2_O at 99 atom% ^15^N enrichment to achieve approximately 5 atom% ^15^N enrichment of N_2_O pool in the chamber headspace, while increasing the chamber headspace gas concentrations by 20 ppb N_2_O and 10 ppb SF_6_. We collected 90 ml of gas samples from the chamber headspace at 0, 15, 30, and 45 min after injection of the spiking gas. The collected gas samples were stored in 60 ml pre-evacuated glass vials sealed with Teflon-coated rubber septa and aluminum crimps and then transported to the laboratory.

**Laboratory assays**

On the same day as soil collection, we performed soil extractions on subsamples of the composited soil samples to characterize soil inorganic N availability and iron (Fe) redox status near the chamber locations. Concentrations of 2M KCl-extractable NH_4_^+^ and NO_3_^-^ were measured on a SmartChem®200 Discrete Chemistry Analyzer (WESTCO Scientific Instruments Inc., France). Concentrations of 0.5N HCl-extractable total Fe and Fe(II) were measured on a Genesys 20 spectrophotometer (Thermo Scientific Spectronic, Waltham, MA, USA) using the ferrozine assay modified from Liptzin & Silver^2^. Iron redox status, as a proxy for the abundance of anaerobic soil microsites, was calculated as the percentage of the total acid-extractable Fe pool accounted for by Fe(II).

Within 24 h of soil collection from the field, we performed a laboratory ^15^N pool dilution and tracer experiment to quantify gross rates of mineralization (GMR) and nitrification (GNR)^3^ and estimate the relative contribution of nitrification and denitrification to soil N_2_O production^4^. Briefly, 300 g of each composited soil sample was split into two Ziploc bags and then held under ambient conditions overnight. The two bags of soil samples received 1 ml of either 99 atom% K^15^NO_3_ or 99 atom% (^15^NH_4_)_2_SO_4_ (Cambridge Isotope Laboratories, Inc., Andover, MA) solution and were mixed thoroughly by hand to homogenize the labeled soil. The concentrations of the labeling solution were determined after the measurement of background NH_4_^+^ and NO_3_^-^ concentrations in the soils with the goal to enrich the existing pool of NH_4_^+^ or NO_3_^-^ to approximate 10 atom%. The initial ^15^N enrichment of the NH_4_^+^ pool averaged 12.2 atom% and ranged 8.6 – 17.4 atom%. The initial ^15^N enrichment for the NO_3_^-^ pool averaged 13.8 atom% and ranged 7.1 – 18.5 atom%. After 15 min, 50 g of soil sample was weighed from each bag and extracted with 150 ml of 2 M KCl. The remaining soil sample in the bag was sealed in a 475-ml Mason jar with a lid fitted with a rubber septum. After incubation at room temperature for 4 h, a 75 ml of gas sample was taken from each jar and stored in a 60 ml pre-evacuated glass vial for CO_2_, N_2_O, and ^15^N_2_O analyses. We determined CO_2_ and N_2_O concentrations on a GC and ^15^N isotopic composition of N_2_O on an IRMS as described above. We calculated the net CO_2_ (a proxy of soil respiration) and N_2_O fluxes from the linear change in concentrations over the incubation period. The ^15^N_2_O produced from soils receiving ^15^NH_4_^+^ and ^15^NO_3_^-^ were assumed to be derived from nitrification and denitrification, respectively^4, 5^. After collection of gas samples, 50 g of soil sample in the jar was extracted for NH_4_^+^ and NO_3_^-^ with 2 M KCl. The extracts were stored in -20℃ prior to the ^15^N isotope analysis by acid-trap diffusion^6^, and the ^15^N isotopic composition was measured on an IsoPrime 100 IRMS coupled to a Vario Micro Cube elemental analyzer (Isoprime Ltd., Cheadle Hulme, UK; Elementar, Hanau, Germany). We calculated GMR and GNR according to Kirkham & Bartholomew^7^ and Hart et al.^3^.

Within 48 h after soil collection, a subsample from each composited soil sample was used to measure concentrations of soil microbial biomass C (MBC) and dissolved organic C (DOC). Soil MBC was determined using direct chloroform extraction of soil subsamples in 0.5 M K_2_SO_4_ as described by Setia et al.^8^. The organic C in the K_2_SO_4_ extracts was quantified on a Shimadzu Total Organic Carbon analyzer (Shimadzu TOC-L-CSH; Shimadzu Corp., Kyoto Japan). We extracted dissolved organic C (DOC) in a 3:1 ratio of deionized water to dry soil equivalent mass, and quantified DOC on a Shimadzu Total Organic Carbon analyzer (Shimadzu TOC-L-CSH; Shimadzu Corp., Kyoto Japan).

Soil pH was measured in a soil slurry consisting of a 1:2.5 mass ratio of air-dried soil and DI water using a pH electrode.

Soil organic C (SOC) and TN concentrations for bulk soil, POC, and MAOC samples were determined on a Vario Micro Cube elemental analyzer (Elementar, Hanau, Germany).

**2. Supplementary Tables**

**Table S1.** Results of repeated-measures ANOVA summarizing soil property differences between consistent nitrous oxide (N_2_O) cold spots and episodic N_2_O hot spots (N_2_O flux class class) on five sampling dates (Date) at the conservation tillage site during the 2021 growing season.

|  | ST | WFPS | Soil pH | Fe redox | NH_4_^+^ | NO_3_^-^ | DOC | MBC | GMR | GNR | SR | POC | MAOC |
| --- | --- | --- | --- | --- | --- | --- | --- | --- | --- | --- | --- | --- | --- |
| Between subjects | | | | | | | | | | | | | |
| N_2_O flux class | 0.92 | 0.43 | 0.74 | 0.83 | 0.87 | **0.03** | **0.02** | 0.48 | 0.37 | 0.95 | 0.61 | **0.003** | 0.38 |
| Within subjects | | | | | | | | | | | | | |
| Date | <0.001 | <0.001 | 0.007 | 0.04 | <0.001 | <0.001 | 0.02 | 0.001 | 0.18 | 0.001 | 0.002 | 0.001 | 0.08 |
| Date × N_2_O flux class | 0.45 | 0.04 | 0.25 | 0.82 | 0.009 | 0.02 | 0.17 | 0.15 | 0.95 | 0.95 | 0.26 | 0.04 | 0.07 |

ST, soil temperature; WFPS, water-filled pore space; NH_4_^+^, soil ammonium concentration; NO_3_^-^, soil nitrate concentration; DOC, water-extractable dissolved organic carbon concentration; MBC, microbial biomass carbon; GMR, gross nitrogen mineralization rate; GNR, gross nitrification rate; SR, soil respiration rate; POC, particulate organic carbon concentration; MAOC, mineral associated organic carbon concentration. Bold *P*-values represent significant difference between consistent N_2_O cold spots and episodic N_2_O hot spots across the five sampling dates.

**Table S2.** Mean (SE) values for soil properties by nitrous oxide (N_2_O) flux class at the conservation tillage site during the 2021 growing season.

|  | May 25 | | | | | June 24 | | | | | July 21 | | | | | August 14 | | | | | October 04 | | |
| --- | --- | --- | --- | --- | --- | --- | --- | --- | --- | --- | --- | --- | --- | --- | --- | --- | --- | --- | --- | --- | --- | --- | --- |
|  | Cold spot | | Hot spot | | Cold spot | | | Hot spot | | Cold spot | | | Hot spot | | Cold spot | | | Hot spot | | Cold spot | | | Hot spot |
| Soil temperature (℃) | 29.6 (0.9) | 29.9 (1.0) | | 24.9 (0.3) | | | 24.4 (0.2) | | 27.8 (0.4) | | | 27.8 (0.5) | | 22.7 (0.7)b | | | 26.7 (0.8)a | | 21.7 (0.6) | | | 22.4 (0.7) | |
| Soil pH | 5.4 (0.1) | 5.6 (0.2) | | 5.6 (0.1) | | | 5.4 (0.1) | | 5.7 (0.2) | | | 5.7 (0.1) | | 5.9 (0.2) | | | 6.1 (0.1) | | 5.8 (0.2) | | | 5.9 (0.06) | |
| NH_4_^+^ (µg N g^-1^ soil) | 2.1 (0.8) | 1.8 (0.6) | | 4.0 (0.9) | | | 3.6 (0.6) | | 1.3 (0.1) | | | 1.7 (0.5) | | 1.0 (0.1) | | | 0.8 (0.1) | | 1.5 (0.2) | | | 1.6 (0.2) | |
| MBC (µg C g^-1^ soil) | 161 (27) | 133 (17) | | 149 (12) | | | 153 (12) | | 89 (25) | | | 145 (39) | | 84 (16) | | | 92 (12) | | 80 (12)b | | | 109 (10)a | |
| GMR  (µg N g^-1^ soil d^-1^) | 5.6 (1.8) | 4.9 (0.9) | | 3.2 (0.8) | | | 2.6 (0.6) | | 2.8 (0.6) | | | 3.0 (0.8) | | 4.2 (0.4) | | | 3.2 (0.5) | | 3.4 (0.7) | | | 2.7 (0.4) | |
| GNR  (µg N g^-1^ soil d^-1^) | 18.9 (5.7) | 16.7 (4.9) | | 10.0 (3.4) | | | 12.9 (3.8) | | 11.4 (3.2) | | | 10.1 (2.2) | | 2.6 (0.6) | | | 3.5 (1.0) | | 6.8 (2.8) | | | 7.2 (2.2) | |
| Soil respiration rate  (µg C g^-1^ soil h^-1^) | 0.74 (0.05) | 0.66 (0.06) | | 0.44 (0.02) | | | 0.48 (0.03) | | 0.46 (0.04) | | | 0.44 (0.03) | | 0.45 (0.04) | | | 0.51 (0.05) | | 0.44 (0.04) | | | 0.49 (0.04) | |
| POC (mg C g^-1^ soil) | 2.0 (0.2)b | 3.1 (0.3)a | | 1.1 (0.4) | | | 1.5 (0.2) | | 1.3 (0.2)b | | | 2.0 (0.2)a | | 1.6 (0.2)b | | | 2.2 (0.3)a | | 1.3 (0.2) | | | 1.7 (0.5) | |
| MAOC  (mg C g^-1^ soil) | 17.5 (1.7) | 19.5 (1.0) | | 16.1 (1.9) | | | 19.5 (1.1) | | 16.4 (1.6) | | | 20.1 (1.2) | | 16.8 (1.8) | | | 20.4 (1.1) | | 16.9 (1.4) | | | 19.8 (0.9) | |

Different lowercase letters within a row indicate statistically significant differences between consistent N_2_O cold spots (n = 7) and episodic N_2_O hot spots (n = 8) at *P* < 0.05 level. NH_4_^+^, soil ammonium concentration; MBC, microbial biomass carbon; GMR, gross rate nitrogen mineralization rate; GNR, gross nitrification rate; POC, particulate organic carbon concentration; MAOC, mineral-associated organic carbon concentration.

**Table S3.** Results of repeated-measures ANOVA summarizing soil property differences between consistent nitrous oxide (N_2_O) cold spots and episodic N_2_O hot spots (N_2_O flux class) on 12 sampling dates (Date) at the conventional tillage site during the 2022 growing season.

|  | ST | WFPS | Soil pH | Fe redox | NH_4_^+^ | NO_3_^-^ | DOC | MBC |
| --- | --- | --- | --- | --- | --- | --- | --- | --- |
| Between subjects | | | | | | | | |
| N_2_O flux class | 0.09 | 0.87 | 0.20 | 0.84 | 0.49 | **<0.001** | **0.005** | **0.02** |
| Within subjects | | | | | | | | |
| Date | <0.001 | 0.03 | <0.001 | 0.14 | 0.17 | 0.22 | 0.003 | 0.20 |
| Date × N_2_O flux class | 0.60 | 0.12 | 0.21 | 0.73 | 0.81 | 0.49 | 0.06 | 0.32 |

ST, soil temperature; WFPS, water-filled pore space; NH_4_^+^, soil ammonium concentration; NO_3_^-^, soil nitrate concentration; DOC, water-extractable dissolved organic carbon concentration; MBC, microbial biomass carbon. Bold *P*-values represent significant difference between consistent N_2_O cold spots and episodic N_2_O hot spots across the 12 sampling dates.

**Table S4**. Mean (SE) soil properties by N_2_O flux class at the conventional tillage site during the 2022 growing season.

|  | Soil temperature (℃) | |  | Soil pH | |  | Soil NH_4_^+^ concentration (µg N g^-1^ soil) | |  | Microbial biomass C  (µg C g^-1^ soil) | |
| --- | --- | --- | --- | --- | --- | --- | --- | --- | --- | --- | --- |
|  | Cold spot | Hot spot |  | Cold spot | Hot spot |  | Cold spot | Hot spot |  | Cold spot | Hot spot |
| May 04 | 14.4 (0.4) | 14.6 (0.3) |  | 6.9 (0.2) | 6.5 (0.1) |  | 46.8 (39.9) | 61.7 (30.0) |  | 31.0 (15.9) | 32.0 (10.8) |
| May 08 | 16.4 (0.2) | 15.8 (0.2) |  | 6.6 (0.1)a | 6.0 (0.1)b |  | 64.8 (42.5) | 38.8 (10.7) |  | 56.6 (9.9) | 65.2 (6.1) |
| May 13 | 27.2 (0.2) | 26.3 (0.5) |  | 6.2 (0.1) | 6.1 (0.1) |  | 33.1 (19.2) | 72.8 (30.3) |  | 64.1 (8.2) | 66.0 (11.0) |
| May 23 | 17.2 (0.4) | 17.3 (0.2) |  | 6.2 (0.1) | 6.0 (0.1) |  | 4.3 (1.7) | 18.9 (9.9) |  | 89.0 (10.1) | 94.1 (10.4) |
| June 02 | 27.2 (0.6) | 26.0 (0.6) |  | 6.2 (0.1) | 6.0 (0.1) |  | 2.0 (0.4) | 7.2 (3.3) |  | 57.4 (4.6) | 56.9 (19.4) |
| June 11 | 24.6 (0.4) | 23.8 (0.4) |  | 6.2 (0.1) | 6.1 (0.1) |  | 1.3 (0.6) | 3.0 (1.5) |  | 61.6 (15.6) | 84.2 (6.0) |
| June 22 | 29.8 (0.6) | 29.6 (0.5) |  | 6.7 (0.2) | 6.8 (0.1) |  | 107 (37.7) | 80.6 (35.9) |  | 71.5 (9.0) | 83.1 (6.6) |
| June 26 | 27.2 (0.2) | 27.6 (0.2) |  | 6.1 (0.1) | 6.0 (0.1) |  | 29.1 (11.8) | 61.8 (30.7) |  | 59.6 (6.5)a | 81.8 (5.4)b |
| July 09 | 27.2 (0.4) | 27.2 (0.3) |  | 6.1 (0.2) | 6.0 (0.1) |  | 6.8 (4.7) | 39.6 (26.7) |  | 32.4 (20.2)b | 75.4 (9.0)a |
| August 05 | 26.2 (0.2) | 26.2 (0.1) |  | 5.9 (0.1) | 6.0 (0.4) |  | 0.8 (0.2) | 20.4 (15.7) |  | 43.8 (4.4)b | 85.3 (11.9)a |
| August 31 | 22.8 (0.2) | 22.8 (0.2) |  | 6.2 (0.2) | 6.0 (0.1) |  | 0.6 (0.1) | 1.8 (1.2) |  | 61.4 (2.8)b | 105.6 (12.4)a |
| October 04 | 15.6 (0.2) | 15.5 (0.3) |  | 6.2 (0.1) | 6.1 (0.1) |  | 0.9 (0.1) | 1.7 (0.9) |  | 81.6 (18.2) | 104.1 (15.6) |

Different lowercase letters within a row indicate statistically significant differences between consistent N_2_O cold spots (n = 5) and episodic N_2_O hot spots (n = 9) at *P* < 0.05 level.

**3. Supplementary Figures**

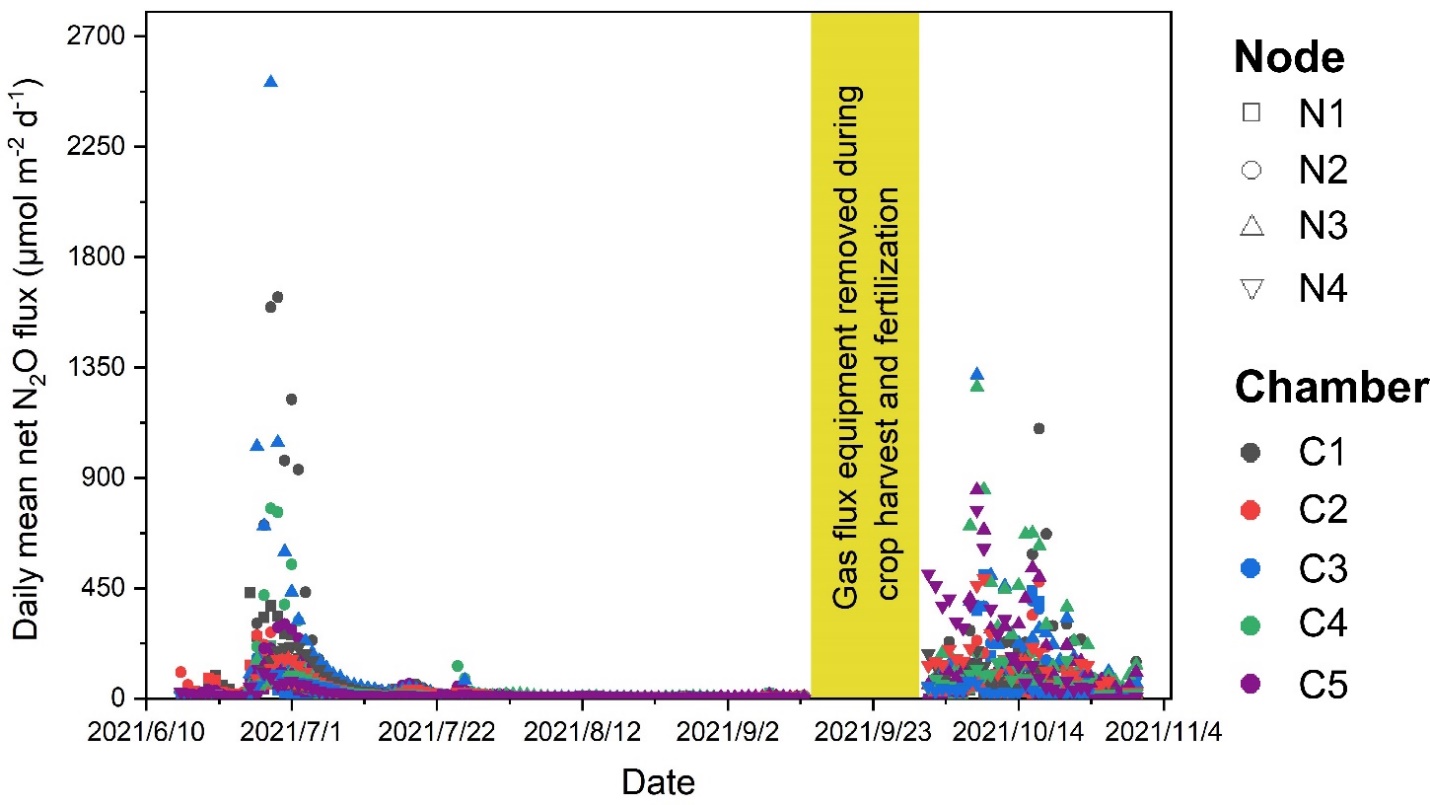

**Figure S1.** Daily average net nitrous oxide (N_2_O) fluxes calculated from hourly flux measurements at 20 autochamber locations at the conservation tillage site during the 2021 growing season. The four sampling nodes (N1-N4) are represented by different symbols, and the five autochambers (C1-C5) at each node are represented by different colors. Spring planting and fertilization occurred on May 01, 2021 and May 7, 2021, respectively. Autochamber measurements began on June 15, 2021, and data through October 31, 2021 were considered used for time stability analysis of growing season N_2_O fluxes.

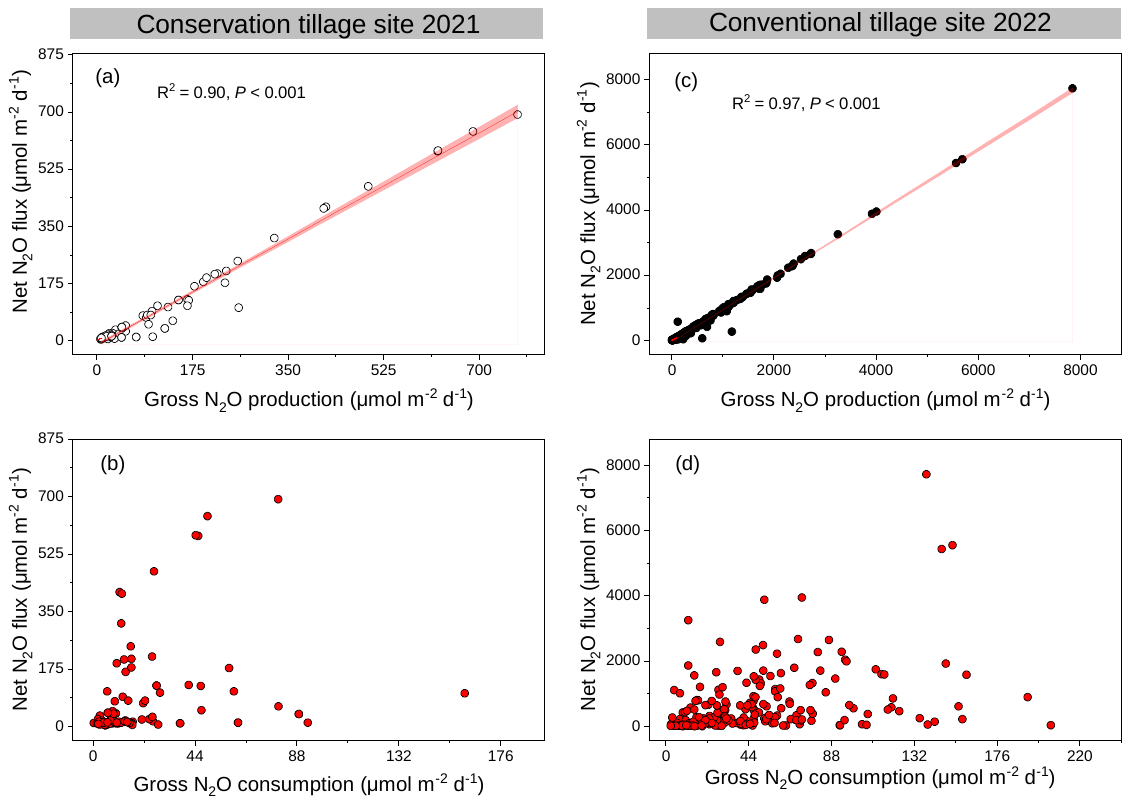

**Figure S2.** Relationships of net nitrous oxide (N_2_O) flux with (a) gross N_2_O production rate and (b) gross N_2_O consumption rate across all five sampling dates and 20 chamber locations (N = 100) at the conservation tillage site during the growing season of 2021. Relationships of net N_2_O flux with (c) gross N_2_O production rate and (d) gross N_2_O consumption rate across all 12 sampling dates and 18 chamber locations (N = 216) at the conventional tillage site during the growing season of 2022. The red lines represent linear regressions with the following equations for panels a and c, respectively: y= 0.93x – 0.16 and y= 0.98x – 0.46.

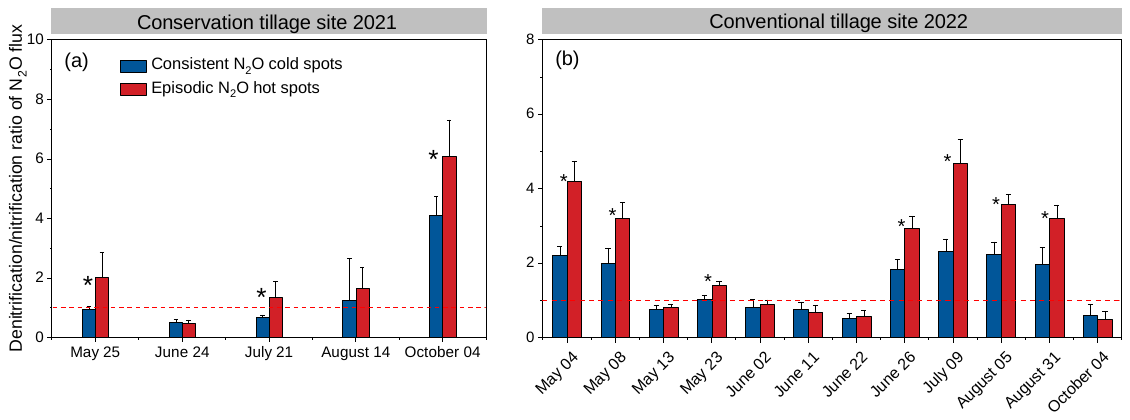

**Figure S3.** The ratio of denitrification- to nitrification-derived net nitrous oxide (N_2_O) flux in the consistent N_2_O cold spots (blue bars) versus the episodic N_2_O hot spots (red bars) measured monthly over the 2021 growing season at the conservation tillage site (a) and measured at 12 time points over the 2022 growing season at the conventional tillage site (b). Ratios above 1 (marked by the dotted line) indicates the dominance of denitrification in N_2_O production over nitrification, and vice versa. Error bars represent standard errors (n=7 and 8 for cold spots and hot spots, respectively, at the conservation tillage site; n=5 and 9 for cold spots and hot spots, respectively, at the conventional tillage site).

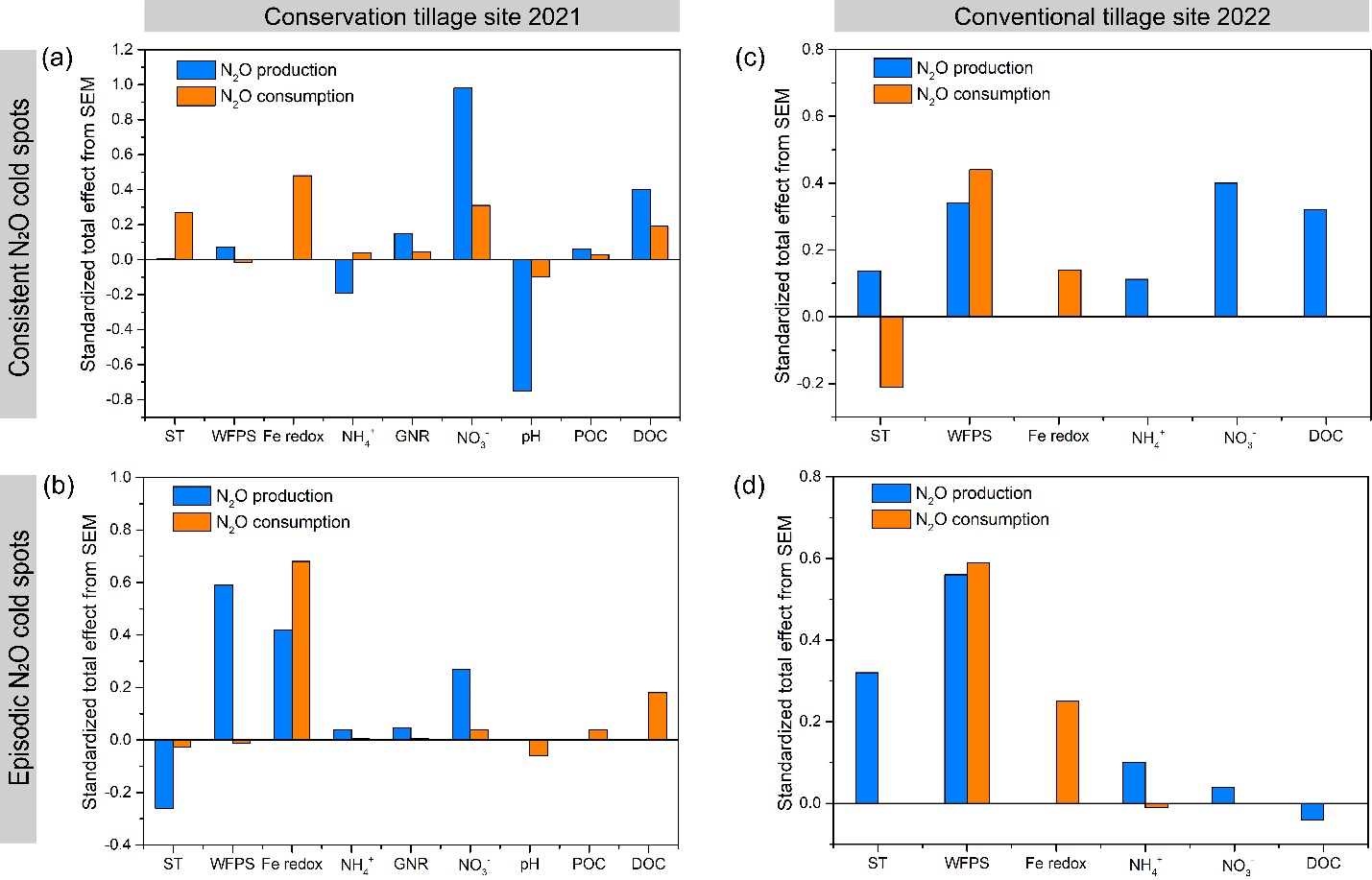

**Figure S4.** Standardized effect from partial least squares structural equation model (PLS-SEM) showing effects of soil variables on gross nitrous oxide (N_2_O) production and consumption in consistent N_2_O cold spots and episodic N_2_O hot spots at the conservation tillage site across five sampling dates during the growing season of 2021 (a and b for cold and hot spots, respectively) and at the conventional tillage site across five sampling dates during the growing season of 2022 (c and d for cold and hot spots, respectively).

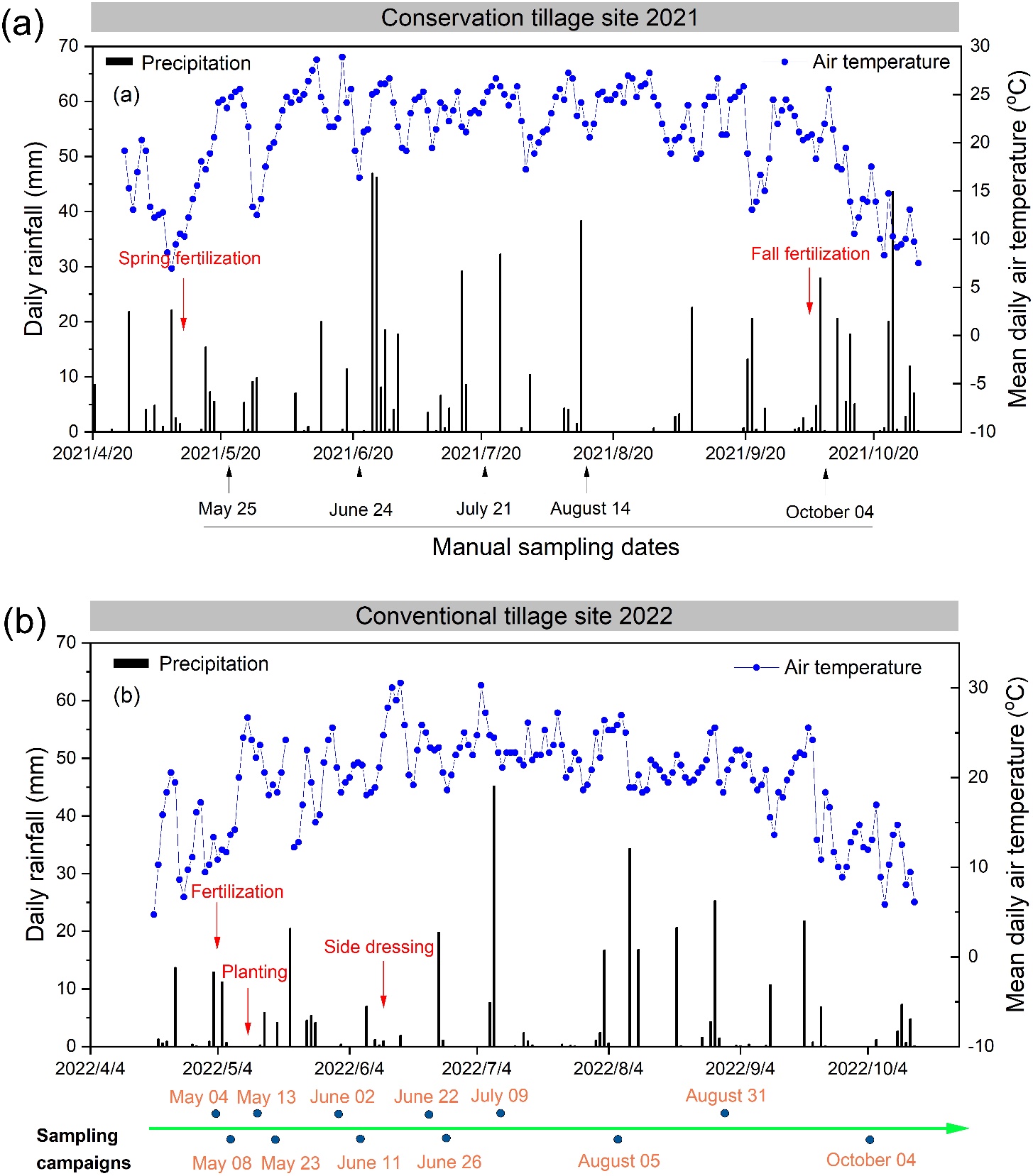

**Figure S5.** Daily rainfall (mm) and mean daily air temperature (℃) at (a) the conservation tillage site and (b) the conventional tillage site during the sampling period in 2021 and 2022, respectively. Field management events and sampling dates are noted.

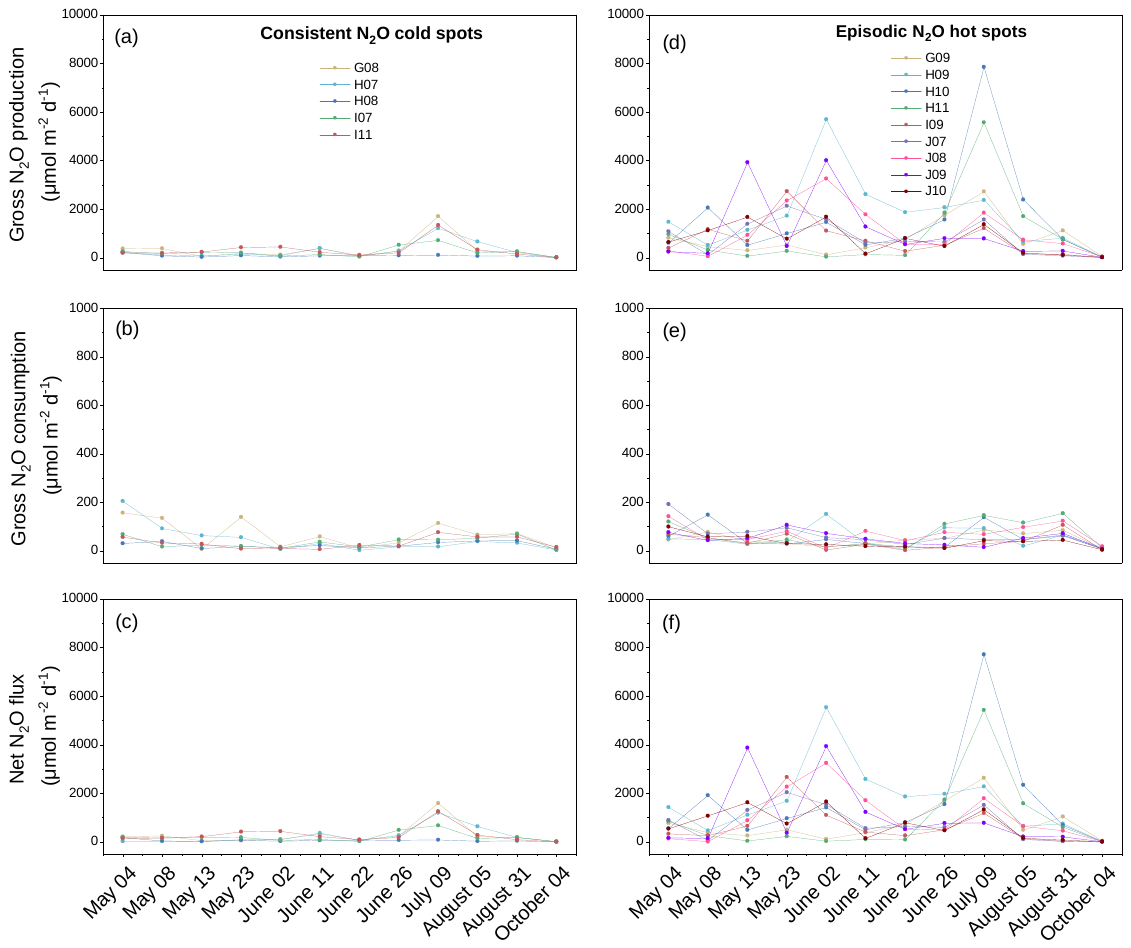

**Figure S6.** Gross nitrous oxide (N_2_O) production rate (a, d), gross N_2_O consumption rate (b, e), and net N_2_O flux (c, f) in consistent N_2_O cold spots and episodic N_2_O hot spots at the conventional tillage site during the growing season of 2022. Colors represent different sampling locations which are identified by grid position.

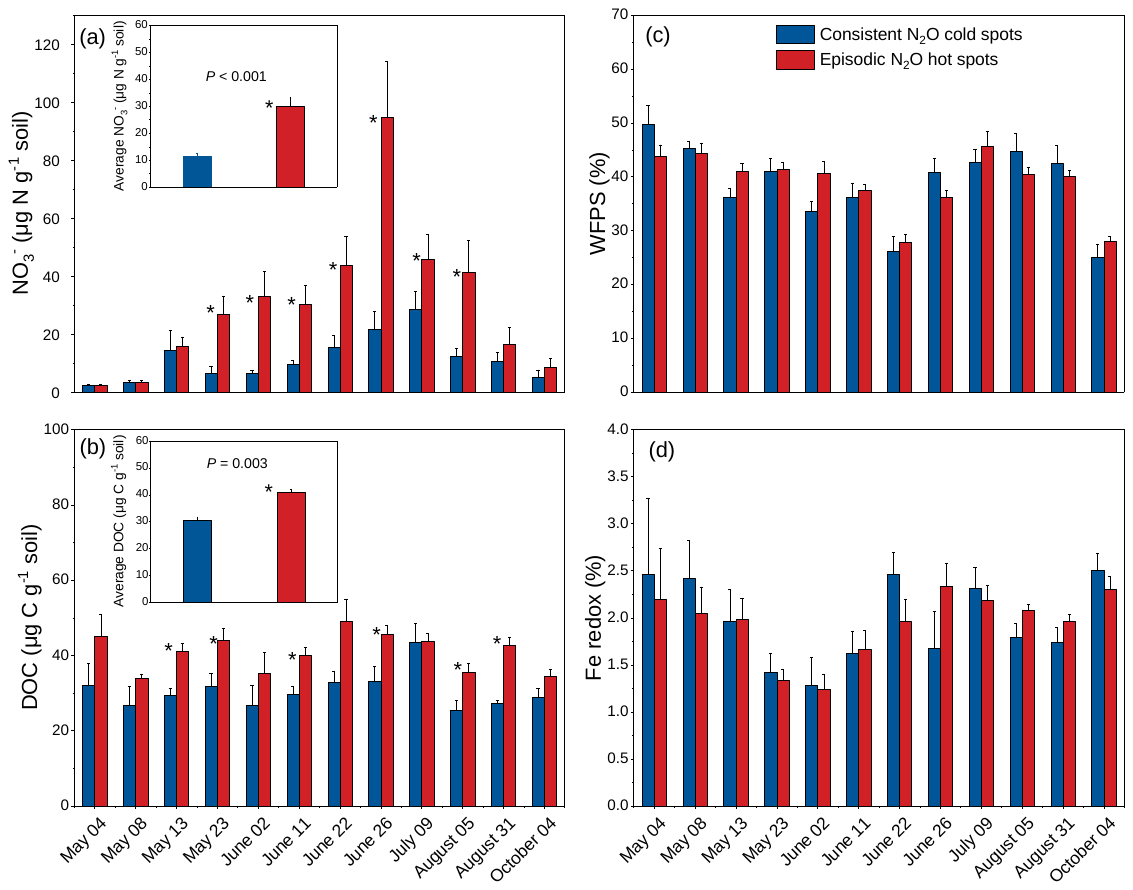

**Figure S7.** (a) Soil nitrate (NO_3_^-^) concentrations, (b) dissolved organic carbon (DOC) concentrations, (c) water-filled pore space (WFPS), and (d) iron (Fe) redox status (Fe(II) percentage of total 0.5 N HCl-extractable Fe pool, an index of anaerobic soil microsites) in consistent nitrous oxide (N_2_O) cold spots (blue bars) versus episodic N_2_O hot spots (red bars) measured at 12 time points over the 2022 growing season at the conventional tillage site. Anhydrous ammonia was injected at a rate of 134.5 kg N ha^-1^ on May 7, 2022, and ammonium thiosulfate and UAN-32 were Y-dropped as side-dress at rates of 13.2 kg N ha^-1^ and 90.2 kg N ha-1, respectively, on June 11, 2022. Planting occurred on May 12, 2022, and harvest occurred on October 28, 2022. Asterisks denote significant differences between cold and hot spots at *P* < 0.05 level. Error bars represent standard errors (n=5 and 9 for cold and hot spots, respectively).

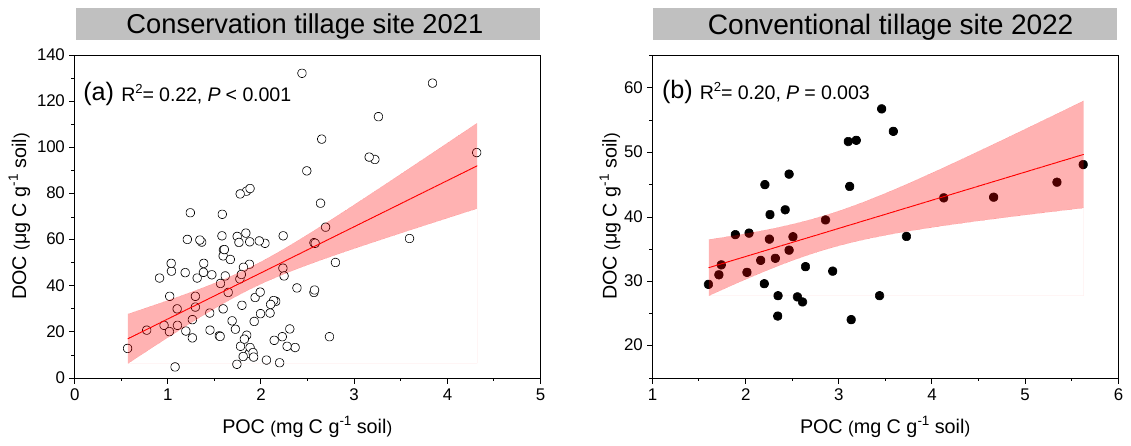

**Figure S8.** Relationship between dissolved organic carbon (DOC) and particulate organic carbon (POC) in soil (a) across all five sampling dates and 20 chamber locations (N =100) at the conservation tillage site during the growing season of 2021 and (b) across two sampling dates (May 13 and August 31) and 18 chamber locations (N =36) at the conventional tillage site during the growing season of 2022. The red lines represent linear regression lines, and the pink shading represents 95% confidence intervals around the regression lines.

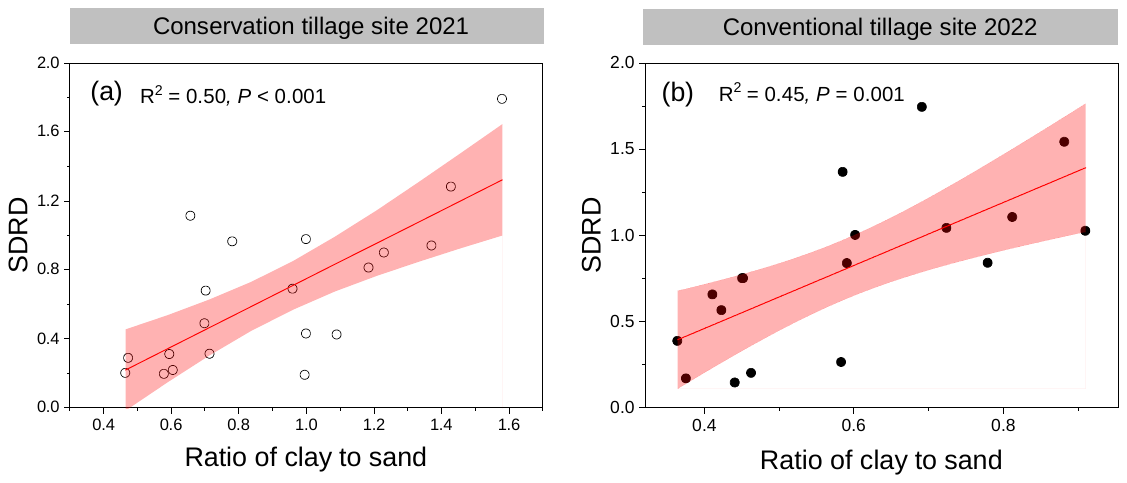

**Figure S9.** Relationship between standard deviation of relative difference (SDRD) of soil net nitrous oxide (N_2_O) emission and the ratio of clay to sand content in soil at (a) the conservation tillage site during the 2021 growing season (N =20) and (b) the conventional tillage site during the 2022 growing season (N =18). The red lines represent linear regression lines, and the pink shading represents 95% confidence intervals around the regression lines.
